## Supplemental Fig. 3 for "Zinc-α2-Glycoprotein Is An Inhibitor Of Amine Oxidase Copper-Containing 3"

Supplemental Fig. 3 of “ZINC-α2-GLYCOPROTEIN IS AN INHIBITOR OF AMINE OXIDASE COPPER-CONTAINING 3”, Matthias Romauch

### **ROS induces ZAG oligomerization**

The difference in the effect of ZAG and LJP1586 on lipolysis at higher octopamine concentrations (Fig. 6) led to the idea that the inhibitory potential of ZAG is coregulated by side-product $H_2O_2$. To test this hypothesis, purified recombinant ZAG was incubated with and without recombinant AOC3, in the presence of $H_2O_2$ or benzylamine (Supplemental Fig. 1, A and B). Incubation with $H_2O_2$ led to a high molecular weight shift (*) in wt ZAG, irrespective of whether AOC3 was present or not. In contrast, addition of benzylamine (and AOC3 dependent production of $H_2O_2$) again led to a high molecular weight shift in wt ZAG, but only in the presence of AOC3. Interestingly, the various glycomutants did not show such behavior. Furthermore, separation of murine plasma by non-reducing SDS-PAGE revealed a very similar WB signal (Supplemental Fig. 1, C). This points to the possibility that $H_2O_2$, whether from AOC3 or not, could induce oligomerization of ZAG *in vitro* and *in vivo*, thereby affecting its biological function.

#### **Figure caption:**

**Supplemental Fig. 1 A and B, WB:** Wt ZAG and its glycomutants were overexpressed in Expi293F cells and affinity purified using their GST-tags. ZAG (after GST removal) was incubated with and without recombinant AOC3, in the presence of $H_2O_2$ (A) or benzylamine (B). Samples were separated by non-reducing SDS-PAGE and proteins were probed using anti-ZAG antibody. **C, WB:** Plasma of a C57Bl6 wt mouse. Proteins were separated by non-reducing SDS-PAGE and proteins were probed using anti-ZAG antibody. Asterisks indicate a high molecular weight protein complex.

### Figures:

### Supplemental Fig. 3

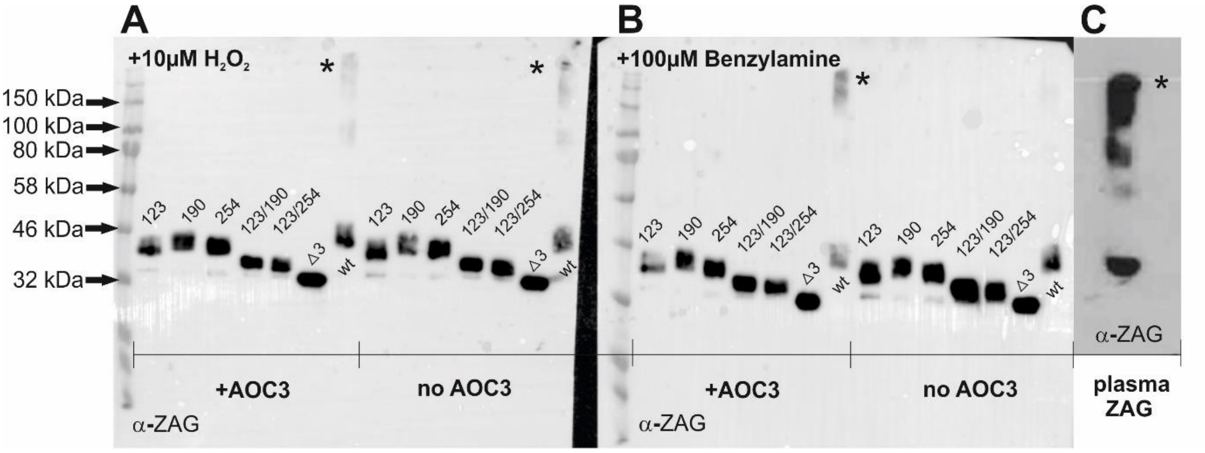
