## Supplemental Fig. 1 and 2 for "Zinc-α2-Glycoprotein Is An Inhibitor Of Amine Oxidase Copper-Containing 3"

Supplemental Fig. 1 and 2 of "ZINC-α2-GLYCOPROTEIN IS AN INHIBITOR OF AMINE OXIDASE COPPER-CONTAINING 3", Matthias Romauch

### Plasma enhances AOC3 activity

Endogenous plasma amine oxidase activity has previously been described in both human and mouse plasma [1–3]. This activity derives from membrane-bound AOC3 which has been released by metalloprotease activity [4]. A pronounced increase in levels of cleaved AOC3 is observed during diabetes, congestive heart failure and liver cirrhosis [5–7]. Incubating recombinant AOC3 with IEX fractions lacking ZAG (Fig. 4, B) increased amine oxidase activity, which might be due to plasma-derived AOC3 activity or other plasma components. To test this hypothesis, the amine oxidase activity in plasma of wt, AOC3 k.o. and ZAG k.o. mice was measured directly using radioactive benzylamine as substrate. The activity linearly increased with measured plasma volume of wt and ZAG k.o. mice, and the activity could be blocked by the highly selective AOC3 inhibitor LJP1586 (Supplemental Fig. 1, A and B). However, AOC3 activity in AOC3 k.o. plasma does not increase with increasing plasma volume and residual activity cannot be further significantly reduced by adding LJP1586 (Supplemental Fig. 1, C). This suggests that AOC3 is the main plasma enzyme responsible for benzylamine deamination, but it does not exclude other amine oxidases that are not sensitive to LJP1586 or that have a higher affinity for other substrates. One such category of enzymes could be the lysyl oxidases, which are also members of the copper amine oxidase family. The lysyl oxidase family is made up of lysyl oxidase (LOX) and the four lysyl oxidase-like enzymes (LOXL1 – 4); these enzymes catalyze the final reaction required for cross-linking of collagens and elastin [8]. Comparison of wt and ZAG k.o. plasma-derived AOC3 activity reveals that lack of ZAG has no influence on activity, which is supported by the fact that the same level of AOC3 protein expression is found in the gonadal white adipose tissue of both wt and ZAG k.o.

mice (Supplemental Fig. 2, D). However, this contradicts the assumption that lack of ZAG automatically leads to significantly higher plasma-derived AOC3 activity.

Since plasma IEX fractions lacking ZAG enhanced recombinant AOC3 activity in a dose-dependent manner (Fig. 4, B), it tested whether plasma in general is able to enhance amine oxidase activity. Indeed, plasma from wt, AOC3 k.o. and ZAG k.o. mice did enhance recombinant AOC3 activity in a dose-dependent manner, reaching maximal activity at 50 µg/ml (Supplemental Fig. 1, D). Since no significant difference among genotypes was observed, it was concluded that a plasma component present in all three genotypes must be responsible. Therefore, bovine serum albumin (BSA; fatty acid-free) was tested and was also found to enhance AOC3 activity in a dose-dependent manner (Supplemental Fig. 2, A). However, combining both plasma (50 µl = 5 mg/ml) and BSA (2 mg/ml) does not further enhance plasma-derived or recombinant AOC3 activity, which indicates that AOC3 activity is already maximally enhanced by the albumin within plasma (Supplemental Fig. 2, B). Interestingly, when amounts of ZAG-IEX fractions and IEX fractions lacking ZAG (Fig. 4, B) were increased, there was no corresponding increase in basal activity (i.e. without addition of recombinant of AOC3) (Supplemental Fig. 2, C), as observed for wt and ZAG k.o. plasma (Supplemental Fig. 1, A and B). This might be due to dilution of plasma-derived AOC3 activity, but also suggests the existence of a non-enzymatic component in IEX fractions lacking ZAG that boosts recombinant AOC3 activity. The latter notion is supported by the work of Dalfo et al., who described a low molecular weight plasma component that, in association with lysophosphatidylcholine (LPC), boosts AOC3 activity by up to 5-fold [9]. This is similar to the effect of incubating recombinant AOC3 with 200 µl IEX fractions lacking ZAG (Fig. 4, B).

### Figure captions:

**Supplemental Fig. 1 A, B and C, [$^{14}$C]-benzylamine assay:** Different volumes of murine wt, AOC3 k.o. and ZAG k.o. plasma were tested for amine oxidase activity. In parallel, the same volumes were tested in the presence of the highly selective AOC3 inhibitor LJP1586. **D, [$^{14}$C]-benzylamine assay:** recombinant AOC3 (50 ng) activity in the presence of plasma of wt, AOC3 k.o and ZAG k.o mice (male, C57Bl/6 genetic background).

**Supplemental Fig. 2 A, [$^{14}$C]-benzylamine assay:** Recombinant AOC3 (50 ng) activity in the presence of BSA (bovine serum albumin; fatty acid-free). **B, [$^{14}$C]-benzylamine assay:** Comparison of AOC3 activity in wt and ZAG k.o. plasma. For basal plasma AOC3 activity (i.e. without recombinant AOC3, but with 50 ng GST), 50 µl (final concentration 5 mg/ml) of plasma with and without BSA (final concentration 2 mg/ml) were tested. In parallel, the same conditions were tested in the presence of AOC3 (50 ng). **C, [$^{14}$C]-benzylamine assay:** AOC3 activity of ZAG-IEX fractions and IEX-fractions lacking ZAG (Fig. 4, B) without addition of recombinant AOC3. AOC3 was replaced with GST (50 ng). **D, WB:** Plasma membrane proteins from gonadal white adipose tissue (1 µg) of wt and ZAG k.o. mice (four per genotype) were separated by SDS-PAGE and probed using α-AOC3 antibody.

**Figures:**

**Supplemental Fig. 1**

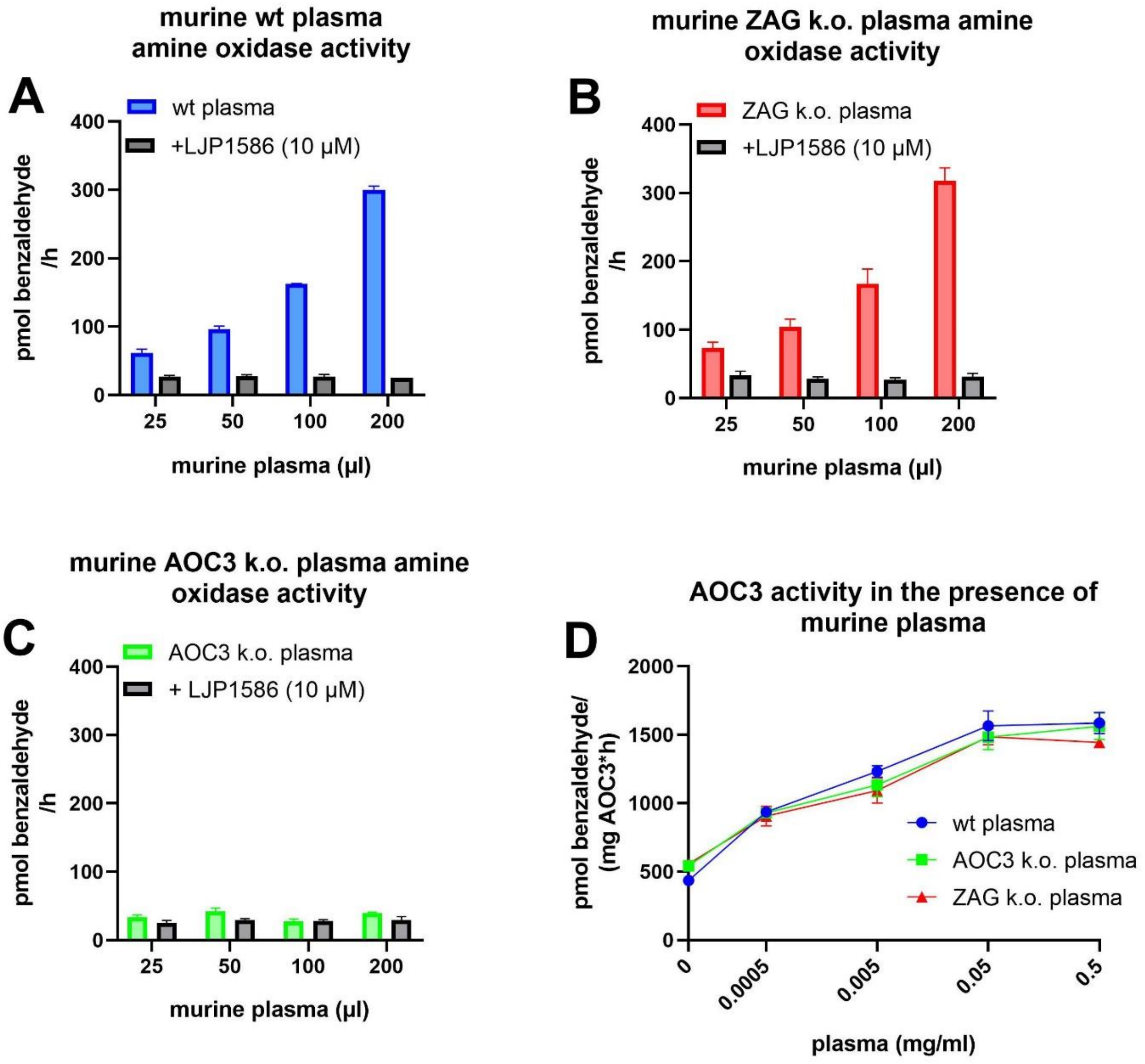

**Supplemental Fig. 2**

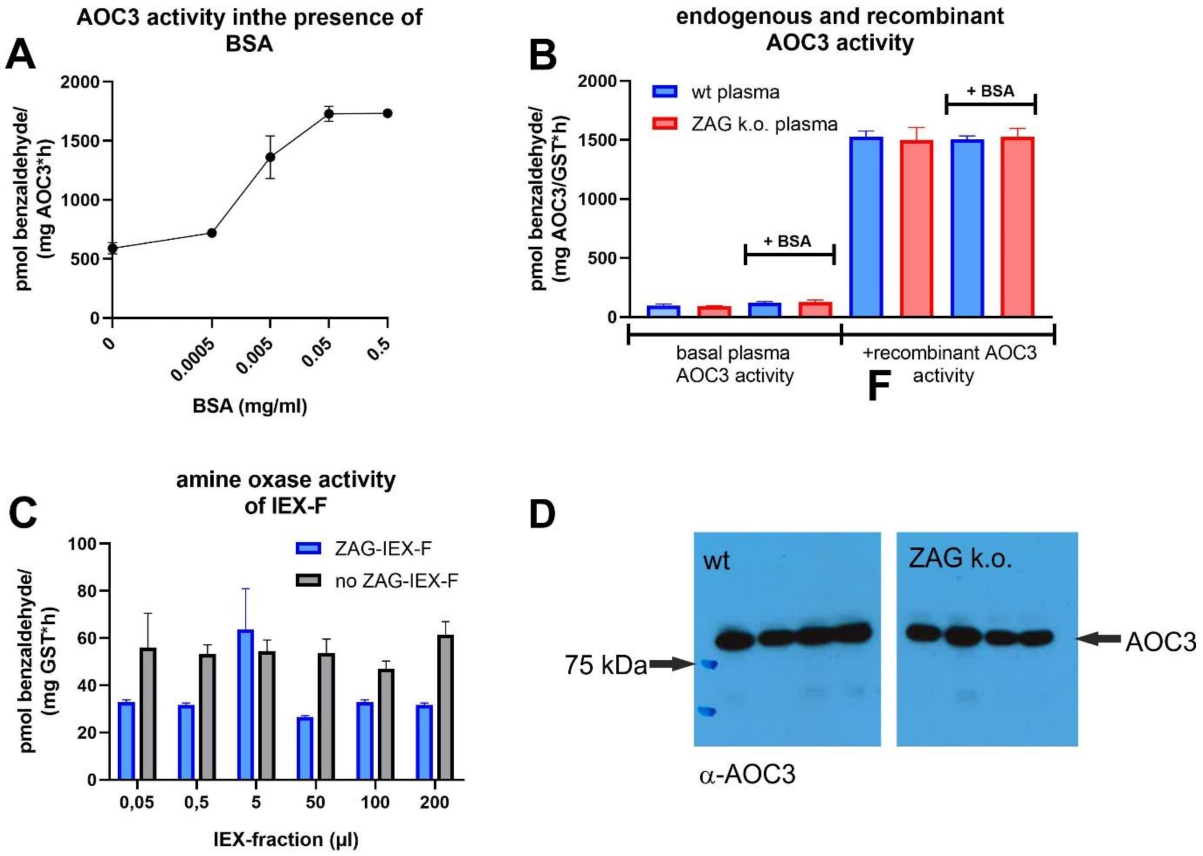
